# Multi-clonal evolution of MDR/XDR *M. tuberculosis* in a high prevalence setting in Papua New Guinea over three decades

**DOI:** 10.1101/172601

**Authors:** Arnold Bainomugisa, Evelyn Lavu, Stenard Hiashiri, Suman Majumdar, Alice Honjepari, Rendi Moke, Paison Dakulala, Grant A. Hill-Cawthorne, Pandey Sushil, Ben J. Marais, Christopher Coulter, Lachlan J. M. Coin

**Affiliations:** Faculty of Medicine, University of Queensland, Brisbane, Australia; Central Public Health Laboratory, Port Moresby, Papua New Guinea; Western Province Health office, Western Province, Papua New Guinea; Burnet Institute, Melbourne, Australia; National Department of Health, Port Moresby, Papua New Guinea; School of Public Health, University of Sydney, Sydney, Australia; Marie Bashir Institute for Infectious diseases and Biosecurity, University of Sydney, Sydney, Australia; Queensland Mycobacterium Reference Laboratory, Pathology Queensland, Brisbane, Australia; Institute for Molecular Biosciences, University of Queensland, Brisbane, Australia

**Author notes:** Contributed equally (Arnold Bainomugisa, Evelyn Lavu, Christopher Coulter, Lachlan J. M. Coin).

## Abstract

An outbreak of multi-drug resistant tuberculosis has been reported on Daru Island, Papua New Guinea. The *Mycobacterium tuberculosis* strains driving this outbreak and the temporal accrual of drug resistance mutations have not been described. We analyzed 100 isolates using whole genome sequencing and found 95 belonged to a single modern Beijing strain cluster. Molecular dating suggested acquisition of streptomycin and isoniazid resistance in the 1960s, with virulence potentially enhanced by a *mycP1* mutation. The outbreak cluster demonstrated a high degree of co-resistance between isoniazid and ethionamide (80/95; 84.2%) attributed to an *inhA* promoter mutation combined with *inhA* and *ndh* coding mutations. Multidrug resistance (MDR), observed in 78/95 samples, emerged with the acquisition of a typical *rpoB* mutation together with a compensatory *rpoC* mutation in the 1980s. There was independent acquisition of fluoroquinolone and aminoglycoside resistance; with evidence of local transmission of extensively-drug resistant (XDR) strains from 2009. These findings underscore the importance of whole-genome sequencing in informing an effective public health response to MDR/XDR *M. tuberculosis.*

## Introduction

Globally, an estimated 10.4 million cases of tuberculosis (TB) occurred in 2015 and *Mycobacterium tuberculosis* was the leading single pathogen killer on the planet (1). The emergence and spread of drug resistant *M. tuberculosis* strains pose a major challenge to global TB control (2). Multi-drug resistant (MDR) TB, which is resistant to at least isoniazid and rifampicin accounted for an estimated 480,000 new cases and 250, 000 deaths in 2015 (1). Additional resistance to fluoroquinolones and second-line injectables defines extensively drug-resistant (XDR) TB (3). In *M. tuberculosis*, drug resistance occurs mainly due to the accumulation of chromosomal resistance conferring mutations without evidence of lateral gene transfer (4). The emergence of drug resistance is dependent on the rate of acquisition of resistance conferring mutations and the frequency with which these drug resistant strains are transmitted within the community (5, 6). Compensatory mutations that limit the fitness cost imposed by drug resistance may enhance the clonal spread of the most successful drug resistant strains (6-8).

Genotyping of *M. tuberculosis* using 24-locus mycobacterial interspersed repetitive unit (MIRU-24) facilitates TB outbreak investigation (9). However, MIRU-24 interrogate small genomic regions that are susceptible to homoplasy and offers sub-optimal discriminatory power (10). The discriminatory power of MIRU-24 is of particular concern with Beijing lineage strains (11). Whole genome sequencing offers optimal resolution to explore local transmission dynamics while also revealing the molecular mechanisms and evolution of drug resistance (6, 12, 13).

Papua New Guinea (PNG) has an estimated TB incidence of 432 per 100,000 population (1). Daru Island, is one of the major hotspots for TB outbreaks and has an estimated incidence rate of 2600 per 100,000 population (14, 15). A recent national MDR TB survey showed that Daru Island had the highest concentration of MDR TB cases in PNG (16) with estimates that 1·5% of the Daru population are diagnosed with MDR TB every year (17) Recent Geo-spatial clustering analysis of TB program data demonstrated evidence of primary transmission on Daru Island (18) but this has not been confirmed by a detailed assessment of molecular epidemiology. The *M. tuberculosis* strains driving the outbreak on Daru Island, as well as the associated drug resistance mutations and temporal accrual of these mutations have not been described. This information is urgently needed to guide patient management and optimal local public health responses.

## Results

### Global phylogeny

We characterized 100 strains from Daru, PNG (Figure 1) by MIRU-24 and whole genome sequencing (Figure 2). MIRU-24 analysis revealed that 95 isolates formed a single dominant outbreak cluster (Figure 3) with low allelic heterogeneity (*h* <0·3) (Table S3). Analysis of SNPs derived from whole genome sequencing revealed that the outbreak cluster formed a monophyletic clade in the Beijing (East-Asian) lineage (modern Beijing sub-lineage), while all the remaining strains belonged to the Euro-American lineage (Figure 4). According to Coll et al classification (19), the outbreak cluster was revealed to be Beijing sub-lineage 2.2.1.1 (Table S4). The Beijing outbreak cluster had a median of 23 differing SNPs between samples (range 0-62), highlighting its limited genetic diversity (Figure S2). The Beijing outbreak cluster was separated by at least 32 SNPs from the nearest neighboring modern Beijing genome included in the representative phylogenetic tree (Table S5).

**Figure 1:**
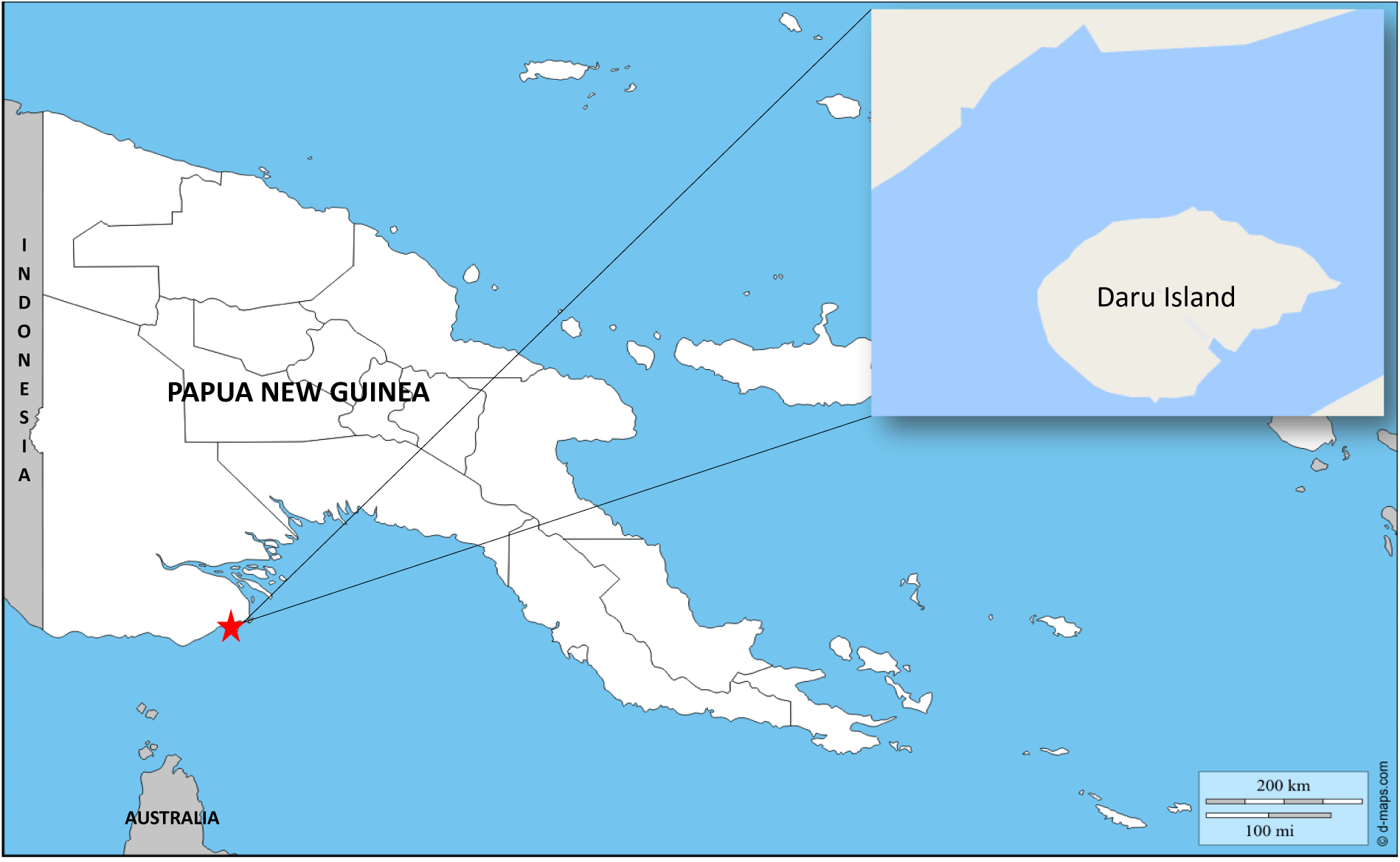
A Map of Papua New Guinea illustrating the study location of Daru Island (Inset) Daru town is the capital of South Fly district, western province of Papua New Guinea. The map was obtained from http://d-maps.com

**Figure 2:**
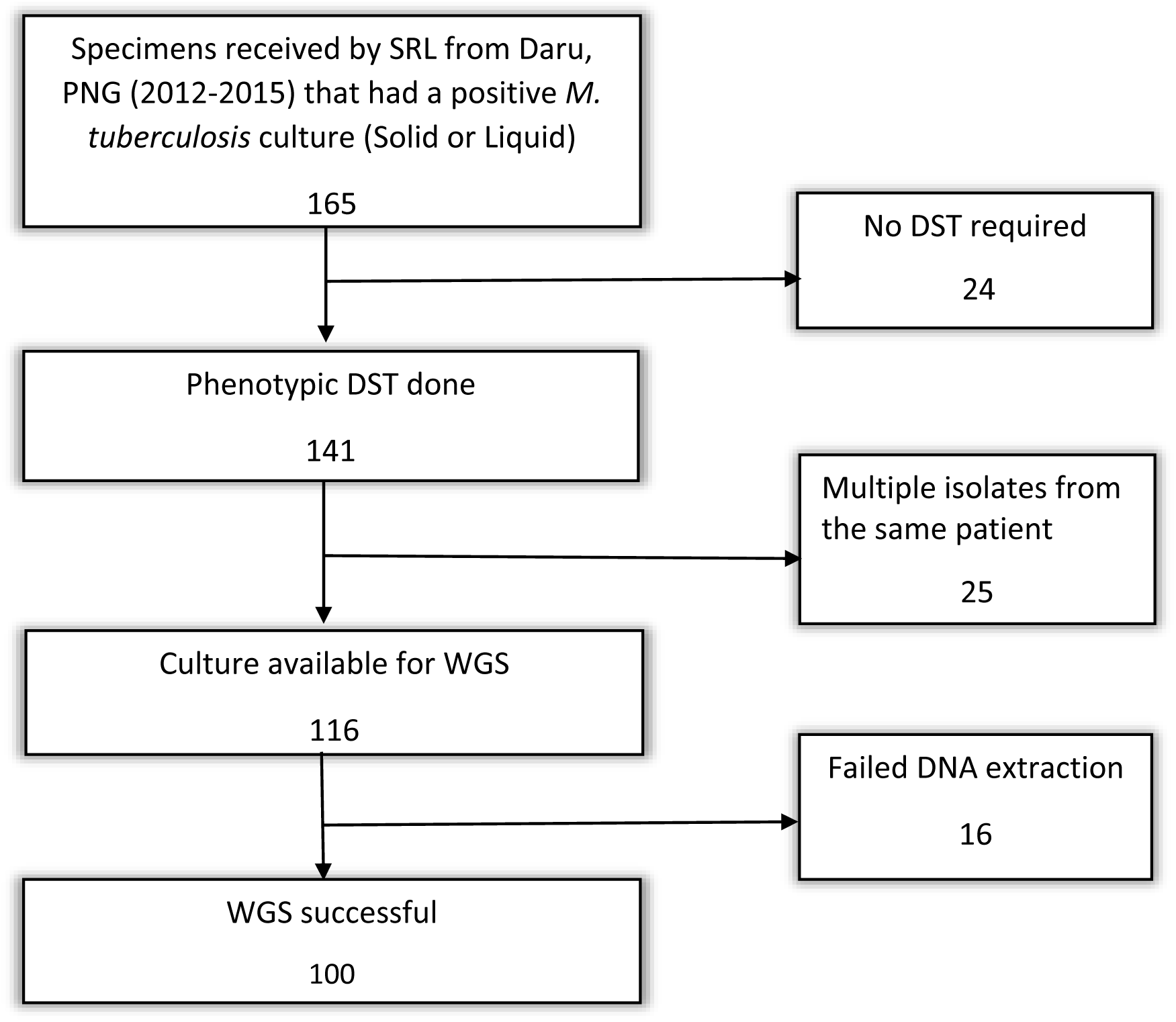
A scheme of workflow to derive clinical isolates included in the study collected between 1/10/2012 to 15/3/2015. PNG – Papua New Guinea; SRL– Supra-National Reference Laboratory; DST – drug susceptibility test; WGS – whole genome sequencing

**Figure 3:**
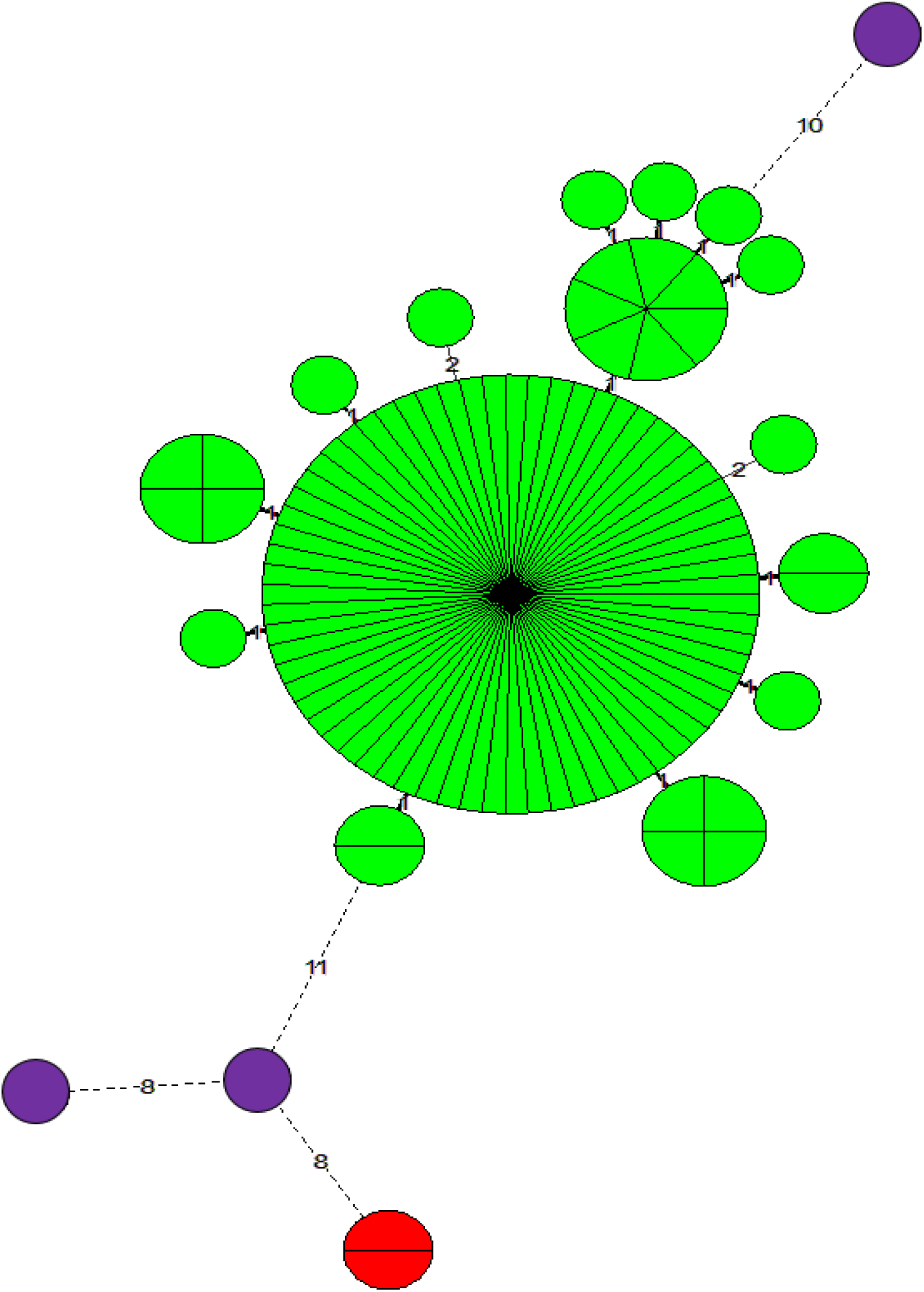
MIRU-24 minimum spanning tree of *M. tuberculosis* isolates collected from Daru Island, Papua New Guinea. Sectioned circles represent two or more isolates that share identical allele profiles. Values on the branch*es* represent allelic difference between isolates. Green-major outbreak cluster, Red-minor cluster, Purple-unique isolates

**Figure 4:**
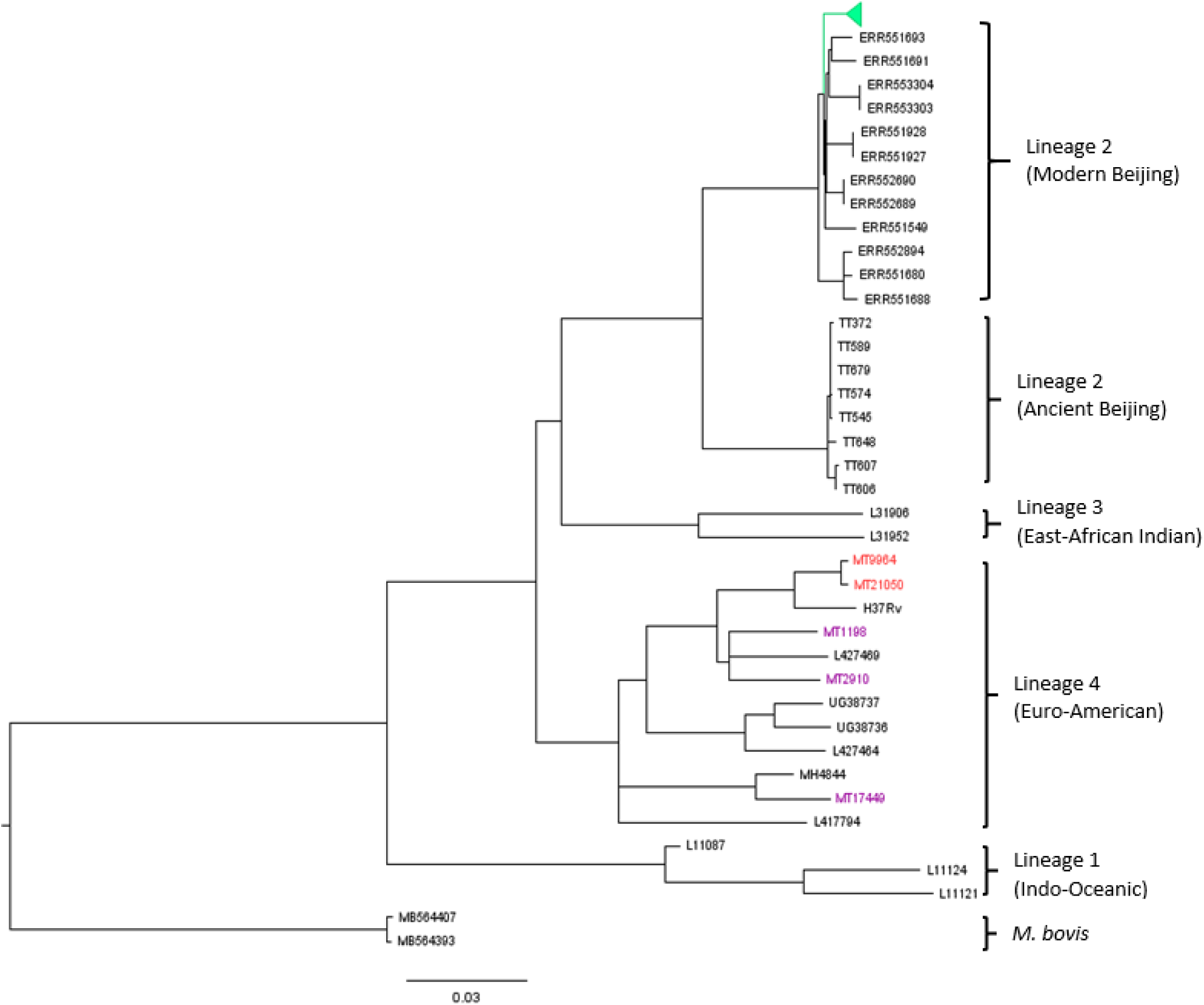
Phylogenetic tree of *M. tuberculosis isolates* collected from Daru Island, Papua New Guinea, together with global representative M. tuberculosis genomes. Tree constructed using 7282 single nucleotide polymorphisms (using RAxML v.7.4.2) and rooted on *M. bovis*. The 95 members of the “outbreak cluster” form a monophyletic clade (green) among the modern Beijing lineage while the colors of the other isolates match those used in Figure 3

### Molecular dating and drug resistance

Table 1 summarizes known and new putative drug resistance mutations detected within the Beijing outbreak cluster, which were parsimoniously mapped to a molecular clock phylogenetic tree that identified four clades; A (n=11), B (n=3), C (n=60) and D (n=21) (Figure 5). Strains in clade A were the most distantly related to the rest of the outbreak cluster and accumulated the least drug resistance mutations. Clade C demonstrated massive clonal expansion and included the majority of the MDR strains (54 MDR and 6 XDR). Ten single nucleotide polymorphisms, including a mutation in *mycP1* (p.Thr238Ala) differentiated clade B, C and D from clade A (Table S6).

**Table 1:**
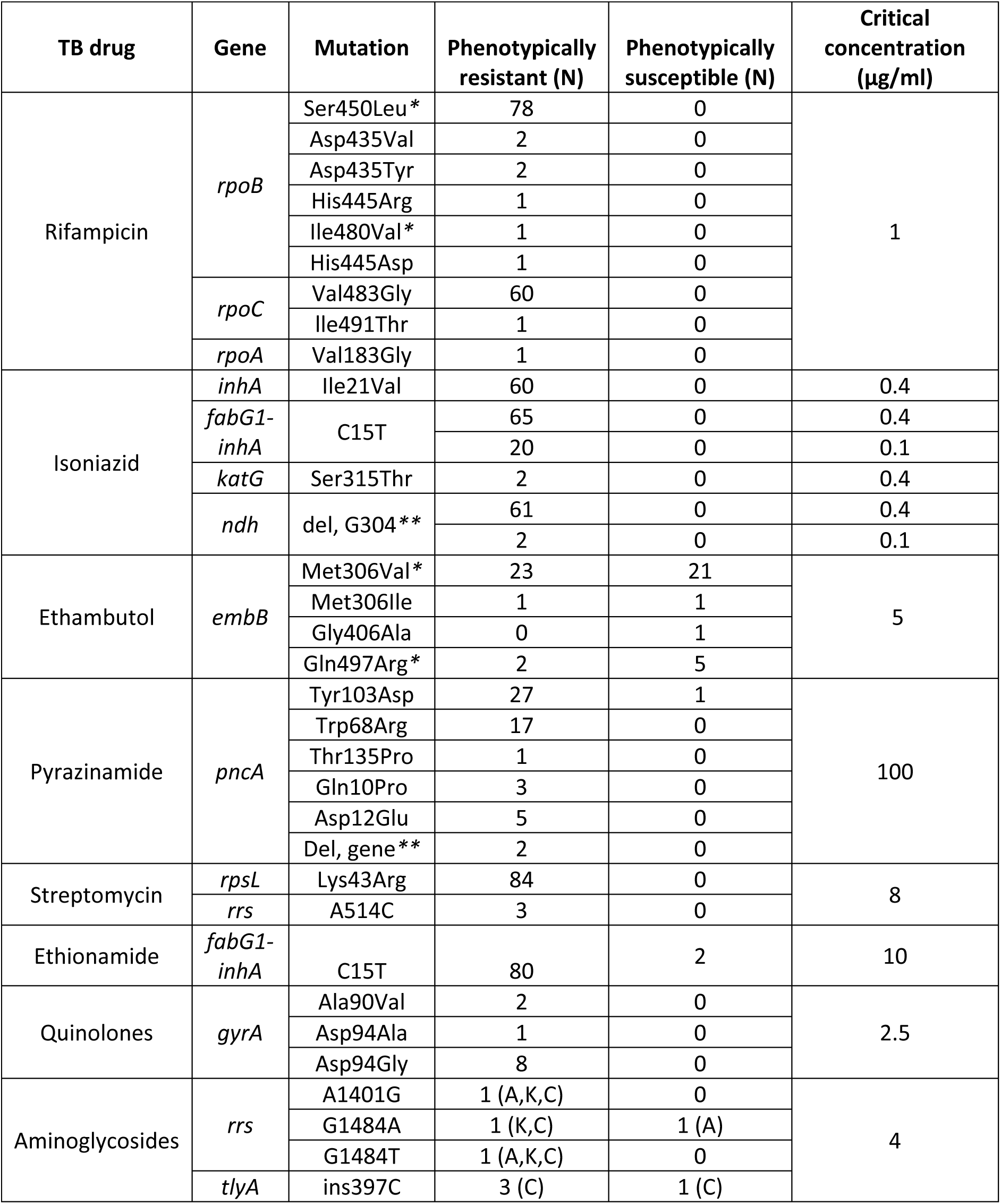
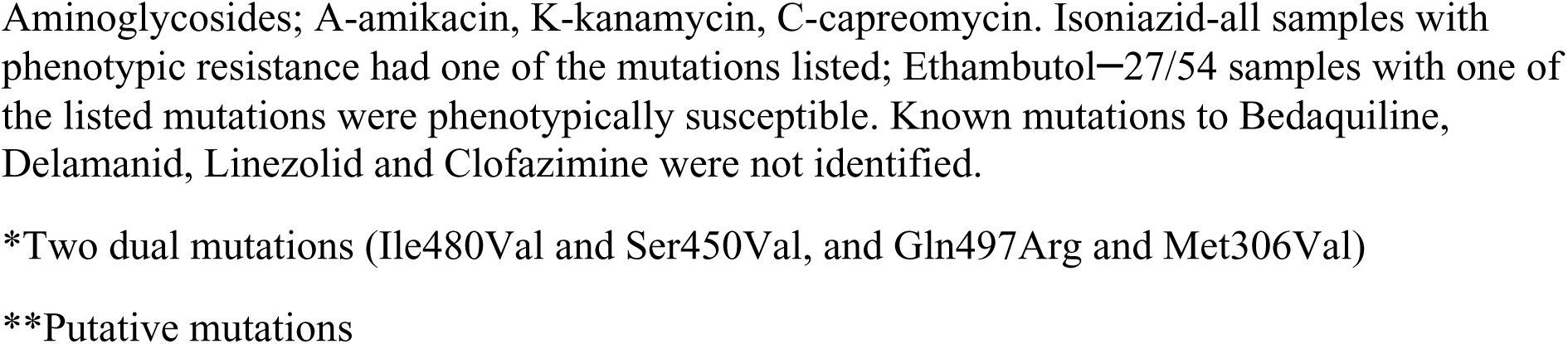
Drug resistance mutations and phenotypic drug resistance observed among Beijing "outbreak cluster" strains

| TB drug | Gene | Mutation | Phenotypically resistant (N) | Phenotypically susceptible (N) | Critical concentration (µg/ml) |
| --- | --- | --- | --- | --- | --- |
| Rifampicin | *rpoB* | Ser450Leu* | 78 | 0 | 1 |
|  |  | Asp435Val | 2 | 0 |  |
|  |  | Asp435Tyr | 2 | 0 |  |
|  |  | His445Arg | 1 | 0 |  |
|  |  | Ile480Val* | 1 | 0 |  |
|  |  | His445Asp | 1 | 0 |  |
|  | *rpoC* | Val483Gly | 60 | 0 |  |
|  |  | Ile491Thr | 1 | 0 |  |
|  | *rpoA* | Val183Gly | 1 | 0 |  |
| Isoniazid | *inhA* | Ile21Val | 60 | 0 | 0.4 |
|  | *fabG1-inhA* | C15T | 65 | 0 | 0.4 |
|  |  |  | 20 | 0 | 0.1 |
|  | *katG* | Ser315Thr | 2 | 0 | 0.4 |
|  | *ndh* | del, G304** | 61 | 0 | 0.4 |
|  |  |  | 2 | 0 | 0.1 |
| Ethambutol | *embB* | Met306Val* | 23 | 21 | 5 |
|  |  | Met306Ile | 1 | 1 |  |
|  |  | Gly406Ala | 0 | 1 |  |
|  |  | Gln497Arg* | 2 | 5 |  |
| Pyrazinamide | *pncA* | Tyr103Asp | 27 | 1 | 100 |
|  |  | Trp68Arg | 17 | 0 |  |
|  |  | Thr135Pro | 1 | 0 |  |
|  |  | Gln10Pro | 3 | 0 |  |
|  |  | Asp12Glu | 5 | 0 |  |
|  |  | Del, gene** | 2 | 0 |  |
| Streptomycin | *rpsL* | Lys43Arg | 84 | 0 | 8 |
|  | *rrs* | A514C | 3 | 0 |  |
| Ethionamide | *fabG1-inhA* | C15T | 80 | 2 | 10 |
| Quinolones | *gyrA* | Ala90Val | 2 | 0 | 2.5 |
|  |  | Asp94Ala | 1 | 0 |  |
|  |  | Asp94Gly | 8 | 0 |  |
| Aminoglycosides | *rrs* | A1401G | 1 (A,K,C) | 0 | 4 |
|  |  | G1484A | 1 (K,C) | 1 (A) |  |
|  |  | G1484T | 1 (A,K,C) | 0 |  |
|  | *tlyA* | ins397C | 3 (C) | 1 (C) |  |

Aminoglycosides; A-amikacin, K-kanamycin, C-capreomycin. Isoniazid-all samples with phenotypic resistance had one of the mutations listed; Ethambutol—27/54 samples with one of the listed mutations were phenotypically susceptible. Known mutations to Bedaquiline, Delamanid, Linezolid and Clofazimine were not identified.
\*Two dual mutations (Ile480Val and Ser450Val, and Gln497Arg and Met306Val)
\*\*Putative mutations

**Figure 5:**
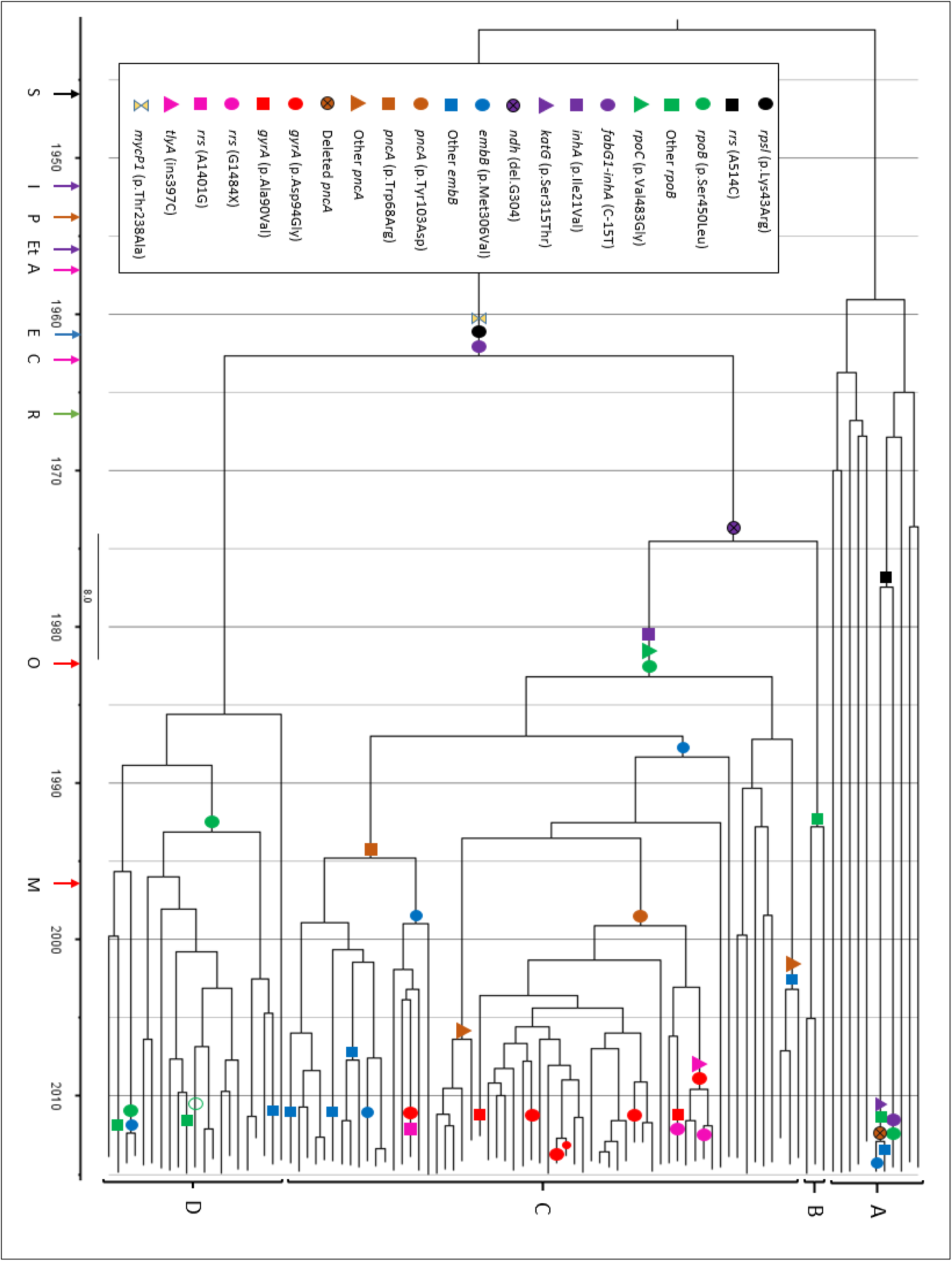
Phylogeny of Beijing outbreak cluster strain with dated acquisition of drug resistance mutations. Four clades (A, B, C and D) were recognizable. Evolutionary gain of resistance conferring mutations and putative mutations for isoniazid (*ndh*) and virulent marker (*mycP1*); parsimonious assigned onto the tree using colored shape with each representing a TB drug or class of TB drug. The colored arrows represent the year of drug discovery; black-S (streptomycin, 1946), green-R (rifampicin, 1966), purple-I (isoniazid, 1952) and Et (ethionamide, 1956), blue-E (ethambutol, 1961), brown-P (pyrazinamide, 1954), red-O (ofloxacin, 1982) and M (moxifloxacin, 1996); pink-A (amikacin, 1957) and C (capreomycin, 1963). Small-red circle-*gyrA* (p.Asp94Ala); hollow-green circle-no *rpoB* mutation

Strains in clades B, C and D demonstrated universal streptomycin resistance conferred by an ancestral *rpsL* (p.Lys43Arg) mutation acquired in the 1960s. The strain ancestral to clade B, C and D also acquired an *inhA* promoter mutation (*fabG1-inhA*, C-15T) associated with low-level isoniazid resistance in the 1960s. The same *inhA* promoter mutation occurred independently in a single clade A strain, while some (3/11) clade A strains acquired streptomycin resistance conferred by an *rrs* (A514C) mutation. Clade C displayed universal high level isoniazid resistance potentially conferred by an intragenic *inhA* mutation (p.Ile21Val) acquired in the 1980s. Occasional high level isoniazid resistance in clade B (1/3) and clade D (2/21) strains were not associated with an *inhA* coding mutation while *katG* (p.Ser315Thr) mutation was detected in only 2 clade A strains.

Of the 87 isoniazid resistant strains tested, 67 had high-level resistance, of which 64 (95·5%) were ethionamide co-resistant; while 16 of remaining 20 (80%) had low-level isoniazid were ethionamide co-resistant. Of the 84 strains tested for ethionamide resistance, 4 strains were susceptible (2 with *katG* mutation but no *inhA* coding or promoter mutations). All strains with co-resistance to at least low-level isoniazid and ethionamide had a *fabG1*-*inhA* mutation. We explored the co-occurring mutations observed in these strains (Table 3). All strains with co-occurring *fabG1*-*inhA*, *inhA* (p.Ile21Val) and *ndh* (del.G304) mutations were ethionamide resistant, while 2/3 with both *fabG1*-*inhA* and *ndh* mutations were ethionamide resistant. No other mutations associated with ethionamide resistance, such as *ethA* or *ethR* were observed. Multi-drug resistance was first acquired by clade C in the 1980s with acquisition of a frequently encountered *rpoB* mutation (p.Ser450Leu), together with a compensatory *rpoC* mutation (p.Val483Gly). The ancestral strain in clade B acquired a different *rpoB* mutation (p.His445Arg) in the 1990s without a compensatory mutation. One rifampicin resistant clade A strain acquired a different *rpoC* compensatory mutation (p.Ile491Thr). The majority of clade D strains (17/21) also acquired the p.Ser450Leu *rpoB* mutation, but at a later time point and only 1 strain had a compensatory *rpoA* (p.Val183Gly) mutation. One rifampicin resistant clade D strain had two *rpoB* mutations (p.Ser450Leu and p.Ile480Val); confirmed by sequence reads spanning both mutations (Figure S3). In a single clade D strain, located in the middle of the *rpoB* (p.Ser450Leu) clone, we could not identify the mutation despite adequate sequence coverage (Figure S4).

There was good correlation between genotypic mutations and phenotypic drug susceptibility determination (Table 1). However, only 50% (27/54) of the strains with putative ethambutol resistance conferring mutations were phenotypically resistant at the critical concentration (5.0 mg/L); MIC testing to encompass lower concentrations of ethambutol was not performed. The *embB* (p.Met306Val) mutation first occurred in clade C during the 1980s with independent acquisition at later time points in clade A, C and D. Another *embB* mutation (p.Gln497Arg) occurred independently on two occasions in clade C.

The majority of clade C (53/60; 88·3%) strains had pyrazinamide resistance mutations. The earliest mutation *pncA* (p.Trp68Arg) was acquired in the 1990s. The only two pyrazinamide resistant clade A strains shared a 3,990bp genomic deletion (position 2287064-2291054, Figure S5) spanning four genes including *pncA*. MDR/XDR strains were more likely to have a *pncA* mutation than non-MDR/XDR strains (56/84 versus 28/84, P < 0.0001).

### XDR TB genotypes

Fluoroquinolone resistance due to mutations in the *gyrA* gene was detected in 11/60 (18%) clade C strains; 6 were XDR TB (Table 1). A single four-member XDR clone emerged around 2009, characterized by an additional capreomycin resistance mutation (*tlyA*, insertion C.397). One XDR strain with phenotypic resistance to amikacin, kanamycin and capreomycin had both *rrs* (G1484T) and *tlyA* (insertion397C) mutations, while another with an *rrs* (G1484A) mutation was phenotypically only resistant to kanamycin and capreomycin. (Figure S6 and Table S7). We assessed the dynamics of drug resistance transmission through enumerating the number of strains that shared known resistance conferring SNPs to infer transmitted versus acquired drug resistance. Primary drug resistance to isoniazid, rifampicin, ethambutol and streptomycin was more dominant in clade B, C and D (P < 0.0001, Table 2). Clade C had additional primary resistance to pyrazinamide and also displayed primary XDR-TB transmission.

**Table 2:** Frequency of clustered (transmitted) drug resistance mutations identified in different clades of the Beijing “outbreak cluster” strains*

|  | Clade A |  | Clade B | Clade C |  | Clade D |  |
| --- | --- | --- | --- | --- | --- | --- | --- |
|  | Independent | Cluster | Cluster | Independent | Cluster | Independent | Cluster |
| *rpoB* | 1 | 2 | 3 | 0 | 60 | 3 | 16 |
| *fabG1-inhA* | 1 | 0 | 3 | 0 | 60 | 0 | 21 |
| *inhA* | 0 | 0 | 0 | 0 | 60 | 0 | 0 |
| *katG* | 0 | 2 | 0 | 0 | 0 | 0 | 0 |
| *rpsL* | 0 | 3 | 3 | 0 | 60 | 0 | 21 |
| *embB* | 2 | 0 | 0 | 3 | 46 | 1 | 2 |
| *pncA* | 0 | 2 | 0 | 0 | 53 | 0 | 0 |
| *gyrA* | 0 | 0 | 0 | 7 | 4 | 0 | 0 |
| *rrs* | 0 | 0 | 0 | 3 | 0 | 0 | 0 |
| *tylA* | 0 | 0 | 0 | 0 | 4 | 0 | 0 |

\*Inferred from figure 5

**Table 3:** Association of isoniazid and ethionamide phenotypes and common genes involved in resistance among Beijing “outbreak cluster” strains

| Gene combinations* | Total | High level<br>INH<br>resistant | Low level<br>INH<br>resistant | Ethionamide<br>resistant | Ethionamide<br>susceptible |
| --- | --- | --- | --- | --- | --- |
| *fabG1-inhA + inhA +*<br>*ndh* | 60 | 60 | 0 | 60 | 0 |
| *fabG1-inhA +ndh* only | 3 | 1 | 2 | 2 | 1<sup>#</sup> |
| *fabG1-inhA* only | 22 | 4 | 18 | 18 | 1 |
| *katG* only | 2 | 2 | 0 | 0 | 2 |
| *inhA* only | 0 | 0 | 0 | 0 | 0 |
| *ethA* only | 0 | 0 | 0 | 0 | 0 |
| *ethr* only | 0 | 0 | 0 | 0 | 0 |

NB: Three strains (*fabG–inhA* only) with low-level isoniazid resistance were not tested against ethionamide as they were not MDR.

## Discussion

Our findings indicate that the large number of MDR TB cases detected on Daru Island in the Western Province of Papua New Guinea is driven by the transmission of a highly drug resistant cluster of modern Beijing lineage strain. Studies from other parts of Papua New Guinea demonstrated a dominance of Euro-American lineage strains, (20, 21) although the contribution of Beijing lineage strains may have been underestimated in these limited surveys. We cannot comment on the geographic spread of this outbreak cluster, but phylogenetic analysis combined with detailed molecular dating suggests that it has been in local circulation since the 1940s and first acquired drug resistance mutations in the 1960s. The identification of four different clades with distinct evolutionary trajectories suggests a ‘permissive’ environment for the *M. tuberculosis* strains to acquire and spread drug resistance within the study setting. Clade C was the most successful clade, acquiring resistance to all four first line drugs and demonstrating clonal spread of both MDR and XDR strains. Phylogenetic analysis indicated that isoniazid and streptomycin resistance was acquired by strains ancestral to clades B, C and D. This is consistent with the use of non-rifampicin containing regimens in the 1960s that were highly reliant on isoniazid and streptomycin (22). Similar analysis done in Russia and South Africa where large MDR outbreaks have been recorded also indicated that streptomycin and isoniazid resistance were acquired before the introduction of rifampicin containing regimens. These settings mostly reported high-level isoniazid resistance due to *katG* mutations (6, 12). In a recent multinational study, *katG* (p.Ser315Thr) was identified as the harbinger mutation that is most frequently associated with future MDR occurrence and risks subsequent clonal spread (23).

Contrary to the experience in most other settings, (23, 24) this Beijing outbreak strain displayed a signature *fabG1*-*inhA* (C-15T) mutation that confers low-level isoniazid resistance with ethionamide co-resistance (25). However, on phenotypic drug susceptibility testing clade C strains demonstrated universal high-level isoniazid resistance. This may be explained by the accumulation of an additional *inhA* (p.Ile21Val) mutation. Such a double mutation may have conferred high-level isoniazid resistance without the fitness cost associated with *katG* mutations, supporting successful clonal expansion and the acquisition of resistance conferring mutations to other anti-tuberculosis drugs (26). An additional consideration is the possibility that the *ndh* frame shift mutation (deletion Glu102fs) observed only in clades B and C may also have contributed, potentially as a compensatory mutation, since SNPs in the *ndh* gene have been associated with increased intracellular NADH/NAD+ ratios hence competitive inhibition of activated isoniazid (27).

The selective advantage of clade C would have been enhanced by the acquisition of a classic *rpoB* (p.Ser450Leu) rifampicin resistance determining mutation together with a compensatory *rpoC* (p.Val483Gly) mutation that would have abrogated its negative fitness effects (7). Similar to previous observations, (28) one strain had two *rpoB* mutations of which p.Ile480Val, which lies outside the 81bp rifampicin resistance determining region, may represent a compensatory mutation. The single strain within the clade D *rpoB* (p.Ser450Leu) cluster, in which we were unable to detect an *rpoB* mutation, may have been from a heteroresistant population since the same *rpoB* mutation was detected in all the neighboring strains. Cultivation of the clinical specimens probably favored selection of the observed strain (29).

An effective host immune response is critical to protect individuals against tuberculosis and to limit ongoing transmission within communities. We identified a non-synonymous *mycp1* (p.Thr238Ala) mutation that is ancestral to clades B, C and D. Mouse models used to identify *M. tuberculosis* ESX-1 substrates and their effects on host cells found *mycP*1 proteins to be essential for early macrophage replication and contribution to virulence by allowing escape of mycobacteria from the phagosome into cytosol of infected macrophages (30, 31). Site directed mutagenesis that inactivated *mycP*1 increased expression of ESAT-6. Altered host immune responses effected by the putative *mycp*1 (p.Thr238Ala) mutation may have increased the virulence and transmissibility of the outbreak cluster since the observed mutation is within the active site (32), but this remains speculative since clade B demonstrated limited clonal expansion.

Previous studies have found a strong correlation between MDR TB and resistance to pyrazinamide or ethambutol, which highlights the problem of drug resistance amplification with continued use of first-line treatment regimens in patients with undiagnosed MDR TB (33, 34). The widespread use of the so-called retreatment regimen or category II posed particular problems, since it added a single drug (streptomycin) to the standard first-line regimen in patients who failed treatment or experienced a second tuberculosis episode. As per World Health Organization (WHO) guidelines, the retreatment regimen was routinely used in the study setting, which would have encouraged amplification of drug resistance given that streptomycin resistance had been present since the 1960s. After a long delay the WHO has finally discouraged the use of the retreatment regimen in its most recent treatment guidance (35).

There was good correlation between genotypic and phenotypic drug resistance profiles, except for ethambutol. This reflects the complex genetic basis of ethambutol resistance, but also emphasizes the highly variable results achieved with phenotypic drug susceptibility testing (13). In our study, phenotypic susceptibility testing for ethambutol demonstrated huge variability, especially when using the Bactec MGIT 960 automated system (36). Safi et al. demonstrated the epistatic nature of ethambutol resistance by identifying that mutations in *embB* are often accompanied by polymorphisms in other genes such as *nuoD* and *Rv3806c* that lead to a progressive increase in the critical concentration (37). We could not identify any of the accompanying mutations previously identified, but hypothesize that a reduction in the critical concentration used may improve the correlation between resistance phenotype and genotype. There is evidence of ongoing transmission of XDR strains characterized by a *tlyA* (ins.C397) mutation that confers capreomycin resistance. One XDR strain demonstrated resistance to all second line injectables, highlighting the ongoing need for timely access to new and repurposed agents like bedaquiline, delamanid and linezolid for effective management of XDR cases. No putative resistance mutations were detected which have been implicated in resistance to these newer agents. Mutations at *rrs* position 1484 may affect aminoglycoside susceptibility differently, since one XDR sample with an *rrs* (G1484T) mutation had phenotypic amikacin resistance, while another with *rrs* (G1484A) mutation was susceptible to amikacin. In a systematic review of studies that assessed the use of second line injectables, the *rrs* (G1484T) mutation was described as a specific predictor of injectable drug resistance but not the *rrs* (G1484A) mutation (38). The Hain MTBDRsl line probe assay recently endorsed by WHO for rapid detection of second-line drug resistance has a probe for *rrs* G1484T but may detect G1484A as a failure of wild type binding (39) and hence incorrectly infer resistance to all second-line injectables; which could mislead treatment.

Our study is limited by its retrospective nature and the fact that clinical isolates tested failed to capture the whole population with MDR TB on Daru Island which is thought to be the nexus of TB transmission. Although this is one of the largest MDR outbreak clusters described to date, the limited number of strains analyzed and the short specimen collection time frame may reduce the accuracy of our time point estimates. However, our estimates have biological plausibility and temporal consistency with the introduction of the respective drugs. Unfortunately a detailed description of when specific drugs were first used in the study setting was unavailable. Finally, we lacked clinical and epidemiological information to enhance our genomic analyses. This information is critical to identify individual and population characteristics that facilitate ongoing transmission of these drug resistant strains.

Whole genome sequencing confirmed a major outbreak of transmitted drug resistant tuberculosis on Daru Island, Papua New Guinea, driven by a single Beijing strain cluster. A major concern is further spread of the outbreak strain to adjacent geographical areas including the capital city, Port Moresby may amplify transmission within Papua New Guinea. There is an urgent need to improve early detection of drug resistant tuberculosis cases with linkage to effective care programs in order to limit drug resistance amplification and terminate ongoing transmission.

## Methods

We performed a retrospective assessment of all clinical isolates referred from Daru Island to the Supra-National Referral Laboratory (SRL) in Brisbane, Australia from October 1, 2012 to March 15, 2015. MIRU-24 and whole genome sequencing were performed to understand the genetic diversity of circulating strains, reveal the mutations associated with drug resistance and to explore the evolution of these mutations. The study was approved by institutional review boards of the University of Queensland, Australia (study number 2015000572) and PNG Medical Research Advisory Committee (study number 16·42).

### Study setting and specimen selection

Daru town located on Daru island (14·7km^2^ in area) is the provincial capital of PNG’s Western Province and has an estimated population of around 15,000 people (40). Daru General Hospital provides health care services to residents of Daru Island and surrounding mainland communities in South Fly District. Clinical specimens are routinely referred to SRL for culture and sensitivity testing. Criteria for referral is on detection of rifampicin resistance by Xpert MTB/RIF assay (Cephid, Sunnyvale, California) on a concurrent sample or at the individual discretion of a clinician.

### Study procedures

DNA was extracted from cultures with confluent growth on Löwenstien-Jensen (LJ) slopes using Spin column High pure PCR template prep kit (Roche Diagnostics, Pennsbury, Germany). MIRU-24 genotyping was performed as per standard protocol (41) and GeneMapper^®^ version 4.0 (Applied Biosystems, California, USA) was used for fragment analysis. Phenotypic drug susceptibility to first-line drugs rifampicin (1·0 µg/ml), isoniazid (0·1 µg/ml; low-level and 0·4 µg/ml; high-level), streptomycin (1·0 µg/ml), ethambutol (5·0 µg/ml) and pyrazinamide (100 µg/ml) was performed based on the proportion method using the automated Bactec Mycobacterial Growth Indicator Tube (MGIT) 960 system (Becton Dickinson, New Jersey, USA). For MDR TB isolates, susceptibility to the second-line drugs amikacin (1·0 µg/ml), capreomycin (2·5 µg/ml), kanamycin (2·5 µg/ml), ethionamide (5·0 µg/ml), ofloxacin (2·0 µg/ml), *p*-aminosalicylic acid (4·0 µg/ml) and cycloserine (50 µg/ml) was determined using MGIT system (20, 42)

DNA for whole genome sequencing was isolated using a lysozyme-phenol chloroform based method and DNA purified using QIAamp DNA mini prep kit (QIAGEN, Hilden, Germany). Paired end libraries were prepared using Illumina Nextera^®^ XT DNA library preparation kit and sequenced using the Illumina MiSeq sequencing platform (San Diego, CA, USA) at Westmead Institute for Medical Research and Australia Genome Research Facility (Sydney, Australia). Raw reads were used to derive octal codes using *insilico* SpolPred (43) and spoligotypes inferred using the international database (SpolDB4) (44). Paired end reads were quality checked using FastQC v0·11·2 (45), trimmomatic v0·27 (46) was used to remove low quality base pairs (Phred score<30) especially at 3’ ends. Trimmed reads were mapped to the *H37Rv* reference genome (GeneBank: NC_000962·3) using BWA-mem with default settings (47). The mean reference coverage was 98·4% (range 96·4-99·8%) and mean high quality base coverage was 70·7X (range 25-182X).

GATK UnifiedGenotyper was used to call SNPs and small indels (48). SNPs and small indels with at least 10X read depth, 80% allele frequency and with at least 10bp difference between neighboring SNPs/indels were retained using customized java and perl scripts. Indels of greater than 4 mutations were excluded from the analysis unless occurring within a known drug resistance associated gene. High quality SNPs and indels were annotated using SnpEff v4·1 (49) and those in repetitive regions such as PE/PPE were excluded from analysis. The SNP allele frequency spectrum was constructed to illustrate the allelic diversity (Figure S1). We characterized mutations in known genes (including regulatory mutations) that confer resistance to rifampicin, isoniazid, ethambutol, streptomycin, pyrazinamide, fluoroquinolones, amikacin, capreomycin, kanamycin, ethionamide, *para*-aminosalicyclic acid, cycloserine, bedaquiline, linezolid and delamanid according to a literature review (Table S1). Raw reads in the form of FASTQ files were submitted to NCBI sequence read archive under the project file number PRJNA385247.

### Allelic diversity, phylogeny and molecular dating

The clonal structure from MIRU-24 was inferred using a minimum spanning tree algorithm implemented in Bionumerics v6·7 (Applied Maths NV, Keistraat 120, Belgium). MIRU-24 profiles with a minimum allele variation of <3 were defined as a cluster. The Hunter-Gaston discriminatory index (HGDI) was used to evaluate the allelic diversity (*h*) of the MIRU-24 locus (50) We obtained 34 global representative genomes from collaborators and from previous studies for phylogenetic analysis (19, 24) Concatenated SNP alignment was used to construct a maximum likelihood phylogenetic tree using RAxML v7·4·2; GTRCAT model at 1000 bootstraps (51) and visualized using FigTree v1·4·2.

Molecular dating of the Beijing outbreak cluster was first performed by Beauti using an alignment containing both invariable and variable sites, the resultant file was used as an input file for BEAST v1·8·2 (52). Base substitution was modelled using Hasegawa-Kishino-Yano (HKY) or General Time Reversible (GTR) model with an estimated base frequency and a gamma distribution among site rate variation with four rate categories. The lognormal relaxed clock (uncorrelated) model which assumes independent mutation rates on different branches, was used (12, 13, 53, 54) The tree was calibrated using time of sample collection as tip dates for each genome, specified in years before the present.

We used uniform prior distribution for all the trees using a mean mutation rate of 0·35 SNP/genome/year (12, 55-57) and compared the model performance under the different demographic models by calculating the Bayes factors from marginal likelihood estimates obtained from path sampling/stepping stone sampling (58). For each analysis, two independent runs of 50 million steps of Markov chain Monte Carlo (MCMC) were performed, discarding 10% burn-in and drawing samples at every 5,000 steps. Three independent runs of the best model were performed for consistency. Tracer was used to examine Markov chain convergence, adequate mixing, chain length and effective sample size (ESS > 200). TreeAnnotator was used to obtain the best supported topology under the maximum clade credibility method.

A constant demographic model that used GTR substitution and gamma distribution at 4 categories was found to be the best fit since it had a better marginal likelihood estimate compared to the other models (Table S2). The overall mutation rate was estimated to be 0·36 SNP/genome/per year (95%High Posterior Density, HPD 0·24-0·48) which is consistent with other published reports (55, 59, 60)

Resistance conferring mutations were parsimoniously mapped on the phylogenetic tree (considering no reversion) and divergence time used to infer the likely timing of drug resistance acquisition. The number of strains that shared known resistance conferring SNPs from the tree nodes (Figure 5) were enumerated to infer primary drug resistance versus non-shared drug resistance conferring SNPs (tree branches) to infer acquired drug resistance (6, 13, 23). A Fisher’s exact test in R statistical package was used to assess the association of observed drug resistance mutations and MDR/XDR phenotypes.

## Author contribution statement

BA, CC, BM, LC, RM, SP, HS, DP, SM, AH and EL designed the study. BA, GC and SP did laboratory work and whole genome sequencing. BA, EL, BM, CC, SP, GC and LC did analysis. All coauthors reviewed and approved the final manuscript

## Competing financial interests

We declare no competing interests

## Acknowledgement

This study was funded by the National Health and Medical Research Council, Australia (grant APP1044986) and AusAid (grant 01 HHISP-2013-0015). We thank the Papua New Guinea National Tuberculosis Program, Provincial Health Staff (Western Province) and staff of Queensland Mycobacterium Reference Laboratory for their assistance. Devika Ganesamoorthy, Derek Benson, Nour Ben Zakour and Alex Outhred for their assistance with data analysis is gratefully acknowledged. Associate Professor Vitali Sintchenko is thanked for his leadership in pursuing WGS strategies in Australia as outlined in NHMRC grant APP1044986. We thank Professor Matthew Cooper for discussions and interpretation of results.

